# Replicating RNA as a component of scrapie fibrils

**DOI:** 10.1101/2023.08.17.553578

**Authors:** Leslie R. Bridges

## Abstract

Recently, electron cryo-microscopy (cryo-EM) maps of fibrils from the brains of mice and hamsters with five infectious scrapie strains have been published^1–5^ and deposited in the electron microscopy data bank (EMDB)^6^. This represents long-awaited near-atomic level structural evidence, widely expected to confirm the protein-only prion hypothesis^7,8^. Instead, the maps reveal a second component, other than protein. The aim of the present study was to identify the nature of this second component, in the published maps^1–5^, using an *in silico* approach. Extra densities (EDs) containing this component were continuous, straight, axial, at right angles to protein rungs and within hydrogen-bonding distance of protein, consistent with a role as guide and support in fibril construction. EDs co-located with strips of basic residues, notably lysines, and formed a conspicuous cladding over parts of the N-terminal lobe of the protein. In one ED, there was evidence of a Y-shaped polymer forming two antiparallel chains, consistent with replicating RNA. Although the protein-only prion hypothesis^7^ is still popular, convincing counter-evidence for an essential role of RNA as a cofactor has amassed in the last 20 years^8^. The present findings go beyond this in providing evidence for RNA as the genetic element of scrapie. To reflect the monotonous nature of the protein interface, it is suggested that the RNA may be a tandem repeat. This is against the protein-only prion hypothesis and in favour of a more orthodox agent, more akin to a virus. Fibrils from brains of patients with Alzheimer’s disease (AD), Parkinson’s disease (PD), amyotrophic lateral sclerosis (ALS) and other neurodegenerations also contain EDs^9^ and may be of a similar aetiology.

---

One of the pedestals of the protein-only prion hypothesis is the idea that scrapie strains are enciphered by protein conformations, rather than nucleic acids^7,10^. On the other hand, it has recently been affirmed that cofactors (including RNA) are essential strain determinants^8^. High-resolution structural models of scrapie fibrils have therefore been eagerly anticipated, in order to resolve this debate. Near-atomic level cryo-EM maps and corresponding protein structures of five scrapie strains^1–5^ have recently been deposited in the EMDB^6^ and protein data bank (PDB)^11^. Whilst there are certainly differences between the protein conformations, there is also a conspicuous second component, within EDs, coordinating with lysines and other basic residues. Lysine-coordinating EDs are a common feature of fibrils from the brains of patients with AD and other neurodegenerations and evidence suggests that they contain RNA^9^. The aim of the present study was to identify the constituent molecule(s) of EDs in scrapie fibrils, by *in silico* methods, using cryo-EM and atomic data from the five scrapie strains^1–5^, from the EMDB^6^ and PDB^11^ public repositories. The results are discussed in the context of the on-going debate about whether scrapie is due to an protein-only prion or a virus-like agent.

## The nature of EDs

Contrary to the protein-only prion hypothesis^7,8^, scrapie fibrils contain a second component. This second component and protein form a novel orthogonal structure, a supported stack (Fig. 1).

**Fig. 1.**
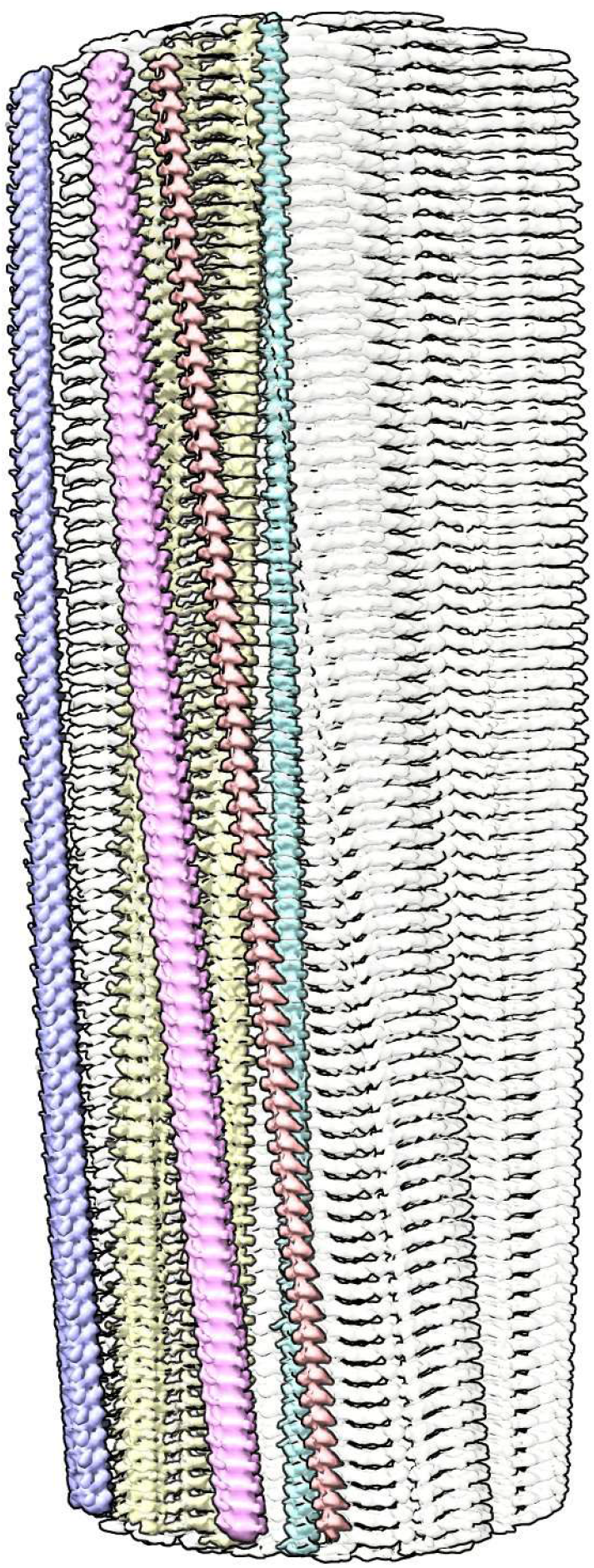
Supported stack. PrP fibril of a22L mouse scrapie. Extra densities (colours) are upright, at right angles to protein rungs (white, transparent), consistent with a structural role as guide and support. Distance between protein rungs is 4.8 Å. Image of EMDB 28089^4^ created with UCSF ChimeraX^21^.

Extra densities (EDs) are present in scrapie fibrils similar to those in Alzheimer’s disease (AD) and other human neurodegenerations^9^. They are continuous with a repeat distance of about 4.8 Å matching that of protein (Fig. 2). They have well-defined features, a 3-blob morphology and Y-shaped connectivity. They are within hydrogen bonding distance of protein and maintain a constant attitude to the protein, twisting and arcing gently with the fibril. They are orthogonal, in the dual sense that they are straight and run at right angles to the protein.

**Fig. 2.**
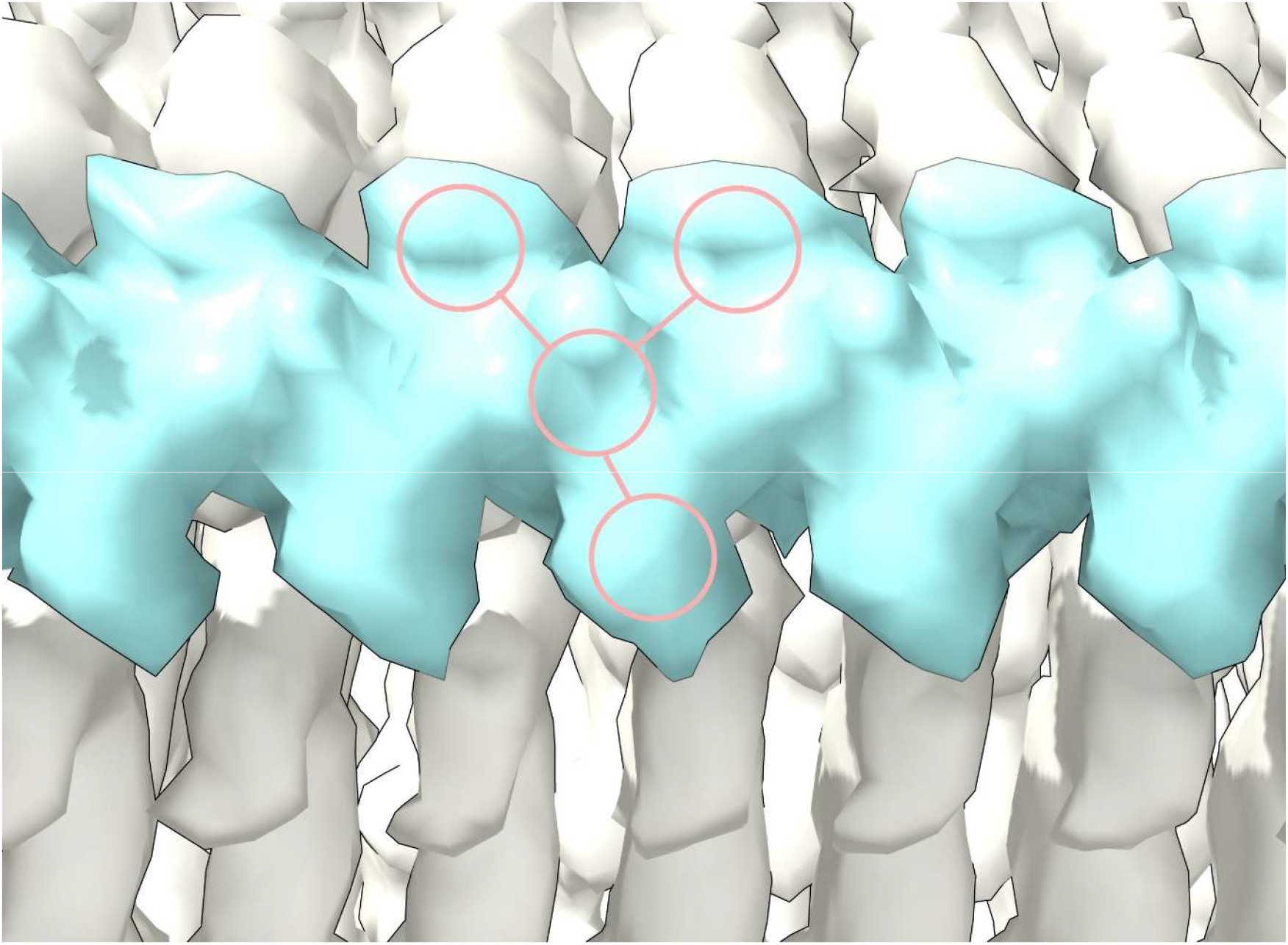
Straight polymer. Extra density (ED) 28089-109 from PrP fibril of a22L mouse scrapie. Note 3-blob pattern and Y-shaped connectivity. The ED is continuous with repeating features consistent with a polymer. It runs at right angles to, in register with, and within hydrogen bonding distance of the protein rungs. The protein rungs (white) are 4.8 Å apart. Image of EMDB 28089^4^ created with UCSF ChimeraX^21^.

The arrangement suggests a supported stack, in which near-planar protein monomers form a stack, supported by uprights composed of the second component. This novel structure, also seen in AD and other human neurodegenerations^9^, is seemingly unique in nature and a hallmark of these diseases.

## The protein environment

Evidence here suggests that the second component is a linear polyanion and a determinant of the protein conformation.

As in AD and other human neurodegenerations^9^, EDs in scrapie coordinate with lysines and other polar residues. EDs in scrapie (Extended Data Fig. 1) coordinate with lysine motifs 100**K**P**SK**103, 103**K**P**K**105, 105**K**T**N**L**K**109 and 109**KH**110 (mouse numbering, residues facing ED in bold). EDs also coordinate with arginines R147, R155 and R163. In some strains, there are contacts with T94, Q159 and Y161 and a floating ED between arginines R155 and R163. In the C-terminal lobe, EDs are found at 184**K**Q**H**186 and 217**Y**Q**K**219 in some strains.

The lysine and arginine patches (K100-K109 and R147-R163) fall within regions highlighted as important in previous research. Residues 90-112 are involved in the conformational transition from PrP^C^ to PrP^Sc^ (refs. 7,12). The central lysine cluster (K100-K109) is predicted to bind charge-neutralising cofactors^13^. Two domains, 91-115 and 144-163, are relatively solvent-protected in infectious (cofactor) PrP^Sc^ compared to non-infectious (protein-only) PrP^Sc^ (ref. 14).

Protein monomers are boomerang-shaped with distinctive angles between lobes (Extended Data Figs. 1 and 2). This angle is determined by variable regions, namely the head of the N arch (residues 112-130) and the central strand (residues 165-175), forming the inter-lobar interface^4,5^.

This is reflected in the root mean square deviation (RMSD) plot comparing a22L and aRML mouse scrapie strains (Extended Data Fig. 2). Since the mice are syngeneic, the differences are unrelated to amino acid sequence. On the other hand, both variable regions are immediately downstream of EDs, suggesting an effect of ligands on protein conformation.

Overall, the evidence suggests that EDs contain a linear polyanion which binds to cationic strips on the surface of the protein and exerts a tethering effect on protein conformation.

## The constituent molecule

Evidence here suggests that RNA is the second component, not merely as a cofactor, but as the replicating genetic material of the scrapie agent.

Considering the fibrils from the Rocky Mountain (RM) lab^1,3,4^, their authors state that EDs may contain non-protein ligands. As exemplar, ED 28089-109 (Extended Data Fig. 3a) is straight (angle 179.5°) with a repeat distance about 4.8 Å matching that of protein. It has a 3-blob morphology and Y-shaped connectivity and its protein environment suggests a polyanion.

A straight form of RNA has been described, complexed to tau fibrils^15^ (Extended Data Fig. 3b). Tau protein was incubated with RNA and formed fibrils in which RNA is straight (angle 179.1°) with a repeat distance about 4.8 Å matching that of protein. It runs parallel to the fibril axis, at right angles to protein rungs and within hydrogen bonding distance of basic residues. The morphology is 3-blob (corresponding to phosphate, ribose and base) and the connectivity is Y-shaped.

Interestingly, Watson and Crick^16^ postulated the existence of a straight form of nucleic acid. Furthermore, straight RNA (named ortho-RNA, oRNA) was modelled, *in silico*^9^, into ED 10650-43 in an alpha-synuclein fibril from multiple system atrophy (MSA)^17^.

1,4-linked polyglucose (Extended Data Fig. 3c), a glycogen-like molecule, is known to form a molecular scaffold in scrapie fibrils^18^, and is rather straight (angle 159.2°) with a repeat distance of 4.6 Å, close to the that of protein. On the other hand, it is single-blob rather than 3-blob, not Y-shaped and not a polyanion.

Polyglutamate (Extended Data Fig. 3d), as beta-strand, is a straight polyanion (angle 179.7°), but multidentate rather than Y-shaped, with a repeat distance of 6.6 Å, out of step with protein, and likely unavailable to brain cells.

Heparin (Extended Data Fig. 3e) is polyanionic but coiled rather than straight (angle 91.1°) with a repeat distance of 12.7 Å out of step with protein. Similar to heparan sulfate^19^, it is likely unavailable inside brain cells.

Poly(ADP-ribose) (Extended Data Fig. 3f) is a potentially bioavailable polyanion^20^ but coiled (angle 126.9°) with a repeat distance of 12.5 Å.

Molecular docking shows feasible poses for oRNA with lysines and arginines, with good AutoDock Vina scores and MolProbity clashscores (Extended Data Fig. 4). The docked poses overlap with EDs and form a rich symmetrical network of hydrogen bonds.

One ED (28089-147), after splitting by proximity to R147 terminal nitrogens, shows two anti-parallel chains arranged as a duplex (Extended Data Fig. 5). The two chains have 3-blob morphology and Y-shaped connectivity and run in opposite directions. They are similar in appearance, and after a 180° rotation placing them in the same direction, can be fitted one into the other (fitmap correlation 0.893 in USCF ChimeraX^21^).

Furthermore, an atomic model of duplex oRNA was posed opposite R147 using molecular docking (Fig. 3). The resulting model has symmetrical hydrogen bonds, good geometry and a reasonable fit to the ED (fitmap correlation 0.767 in USCF ChimeraX^21^). This indicates that duplex RNA is a feasible constituent of the ED. Although uracils are shown in Fig. 3, the duplex structure permits any base sequence in both strands, including any complementary sequences^9^. Thus, whilst the duplex lacks the structure of double-stranded RNA (dsRNA), it could be a refolded form of dsRNA.

**Fig. 3.**
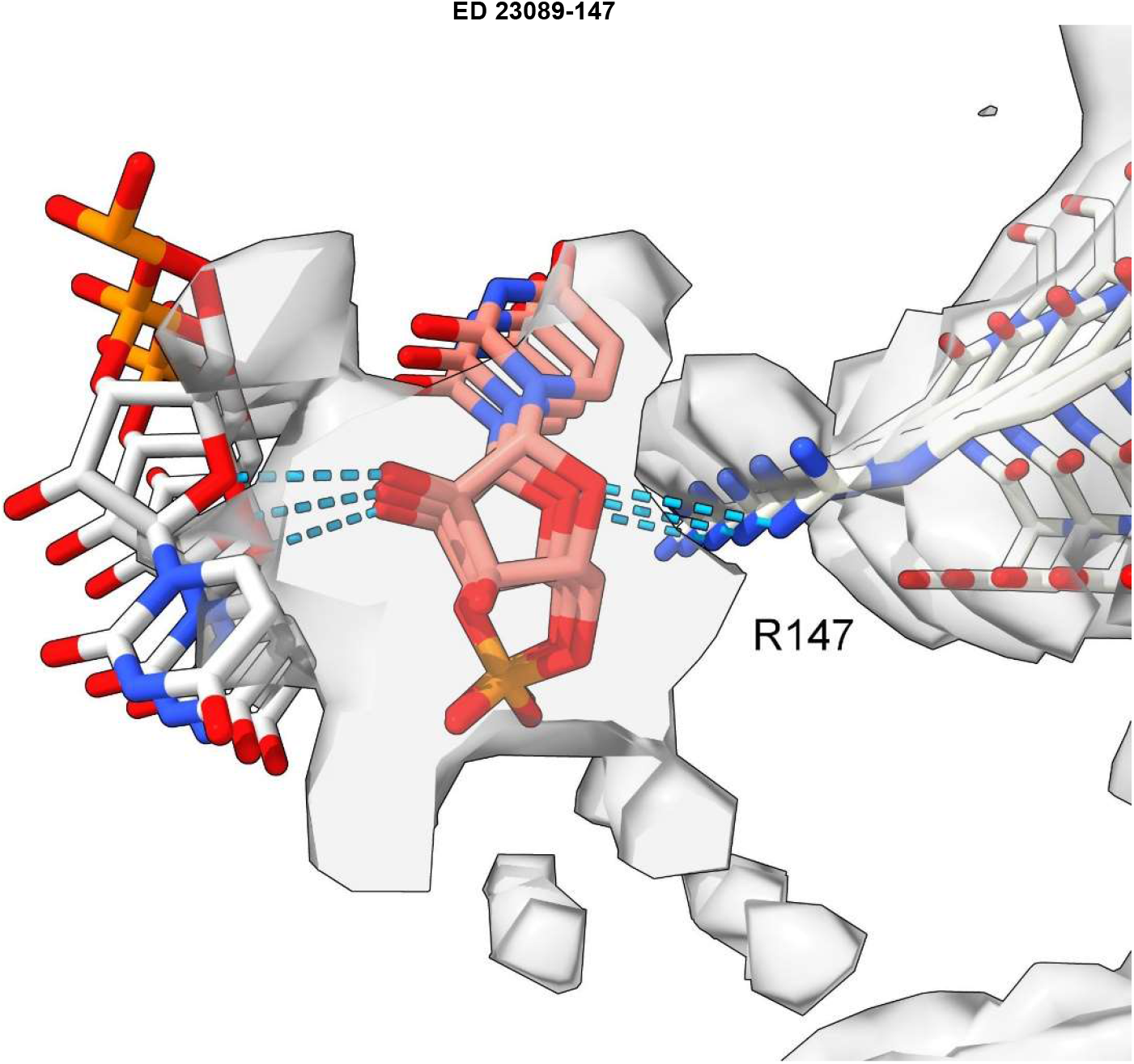
Model of duplex RNA docked to protein. ED 28089-147 from PrP fibril of a22L mouse scrapie. A duplex form of ortho-RNA is docked to protein: good geometry, symmetrical hydrogen bonds and a reasonable fit to the ED are found. The oRNA chains are anti-parallel and congruent with the split ED (see Extended Data Fig. 5). Image of EMDB 28089 and PDB 8efu^4^ created with UCSF Chimerax^21^, oRNA poses generated with AutoDock Vina^40^.

Estimates of molecular weight (MW) were carried out on EDs in the PrP fibril of a22L mouse scrapie. MW per unit of ED were 216 Daltons (Da) for ED 28089-109, 350 Da for ED 28089-147, 389 Da for ED 28089-155 and 94 Da for ED 28089-163. The fused mass of ED overlying K100 to K105 had a MW of 1,029 Da. Such estimations are indeterminate because the occupancy of the constituent molecules is unknown, but EDs at K109 and R163 are consistent with single RNA strands at 67% and 29% occupancies respectively. EDs at R147 and R155 are consistent with duplex RNA at 55% and 61% occupancies respectively. The conglomerate ED between K103 and K105 has a MW equivalent to 6 RNA strands at 54% occupancy.

Other evidence also points to RNA, *viz.* RNA is complexed to PrP in amyloid plaques in scrapie^22^, EM micrographs of scrapie fibrils show tails, sensitive to Zn^2+^ ions, suggestive of RNA^23^ and RNA promotes PrP^Sc^ formation *in vitro*^24^.

Due to averaging, it is not possible to read the base sequences in the present studies. However, given that the protein interface is identical at every rung (Extended Data Fig. 6), for consistent binding the RNA might also be repetitive, either as a homopolymer or short sequence repeat. Host-like repetitive nucleic acid^25^ and single-stranded DNA (ssDNA) with the palindromic sequence (TACGTA)_n_^26^ were found in hamster scrapie. Molecular docking studies show that ssDNA and RNA are potentially interchangeable^9^.

Considering the fibrils from the University College London (UCL) lab^2,5^, the authors assert that EDs represent phosphotungstate (PTA), a polyanionic reagent used in the preparation of their fibrils and not used by the RM lab. The regular appearances are consistent with this idea. However, the presence of EDs in fibrils from both labs suggests that natural ligands are usually present and might therefore have been at least partially displaced in the UCL fibrils.

Overall, the evidence here of paired anti-parallel chains, suggests that RNA is present in scrapie fibrils as the genetic material and not merely a cofactor.

## Comparison with human neurodegenerations

Evidence here suggests that scrapie and human neurodegenerative diseases are part of a spectrum, differing in degree not in kind, and that RNA is a unifying factor.

Features in common include slow tempo and cell-to-cell spread within the brain. Experimentally, they can usually be transmitted to other organisms, although natural transmission is probably confined to certain PrP diseases (notably kuru and scrapie).

Gerstmann-Sträussler-Scheinker disease (GSS) fibrils have two protofilaments whereas scrapie typically only has one (Extended Data Fig. 7). However, core size is smaller in GSS than in scrapie (Extended Data Fig. 8). Whilst the lysine patch (K100-K109) is present in both, EDs are fewer, smaller and less closely-packed in GSS. Furthermore, the arginine patch (R147-R163), known to be associated with infectivity in scrapie^14^, is absent in GSS. The F198S (Indiana kindred) form of GSS^27^ has a typically slow tempo and little or no transmissibility. Furthermore, it has a conspicuous co-pathology, in the form of tau paired helical filaments (PHFs). Arguably, it resembles AD more than scrapie or Creutzfeldt-Jakob disease (CJD), despite being a disease of PrP.

The core sizes of the five scrapie strains range from 132 to 138 residues. In comparison, in human neurodegenerations, core sizes range from 34 residues (beta-amyloid fibril in AD, PDB 7q4b^28^) to 135 residues (TMEM106B fibril in AD, PDB 7qvc^29^). Protofilament sizes of 94 residues (tau fibril in Pick’s disease, PDB 6gx5^30^), 110 residues (tau fibril in progressive supranuclear palsy (PSP), PDB 7p65^31^) and 115 residues (tau fibril in argyrophilic grain disease (AGD), PDB 7p6d^31^) are only somewhat smaller than scrapie. Thus, whilst core sizes are generally smaller in human neurodegenerations compared to scrapie, this is not an absolute distinction. Furthermore, where protofilaments are typically paired, the combined fibril size in terms of residues is, in some instances even larger than scrapie, such as 146 residues (tau PHF fibril in AD, PDB 5o3l^32^). Since the protofibrils in GSS are also paired, the combined residue count is 124 residues (PrP fibrils in GSS, PDB 7umq and PDB 7un5^27^), close to that of scrapie. The possession of key motifs, including lyines, might be more important than core size *per se* in determining infectivity.

Measurements support the visual impression that scrapie fibrils are particularly well endowed with EDs. For the fibrils shown in Extended Data Fig. 9, the volume of EDs compared to the volume of protein density is 13.5% for a22L mouse scrapie (PrP fibril, single protofilament, EMDB 28089, PDB 8efu^4^), 8.5% for GSS (PrP fibril, paired protofilaments, EMDB 26613, PDB 7un5^27^), 9.0% for AD (tau PHF fibril, paired protofilaments, EMDB 26663, PDB 7upe^33^) and 4.3% for MSA (alpha-synuclein fibril, paired protofilaments, EMDB 10650, PDB 6xyo^17^). This extra bulk of EDs could explain the enhanced transmissibility of scrapie, if (as proposed here) it represents replicating RNA. The particular RNA sequence, including palindromicity (see next section), is likely also important.

## The scrapie agent

Based on evidence herein, the following proposals are made about the scrapie agent:

1. RNA is present and forms a unique orthogonal structure with protein.
2. RNA is the genetic element and determines protein conformation.
3. Similar agents cause AD and other human neurodegenerations^9^.
4. The findings are incompatible with the prion hypothesis^7,8^.
5. The agent is also not a virus and may have an earlier lineage. Unlike a virus where RNA and protein are colinear and related by sequence, RNA and protein in the scrapie agent are non-colinear and related by shape.
6. To fully define a scrapie strain, it is necessary to specify both RNA and protein sequences (due to non-colinearity, knowledge of one is not predictive of the other).
7. The agent faces in two directions: its protein-RNA relationship is shape-based but its RNA-RNA relationship is sequence-based (the two replicating strands are Watson-Crick complementary sequences). Its genome is likely a short sequence repeat with no other function than to form more agent with protein.

The shape-based system likely lacks the exclusivity of sequence-based systems such as the DNA double helix. For example, *in vitro*, nucleic acids stimulate PrP_res_ amplification in the rank order poly(A) = poly(dT) > poly(dA) > poly(C)^24^, suggesting non-exclusive, preferential binding. Nevertheless, whilst several RNAs might promote fibril formation, in the life cycle of the agent as a whole, kinetic and thermodynamic factors may narrow this down to one or a few truly effective RNA partners.

The RNA-protein shape paradigm suggests a ready explanation of the species barrier. If a new host protein is able to form an equivalent fold, it will stack with the inoculated fibril and the RNA will stay the same. If it partially aligns, the fibril will change shape and a new RNA will be selected (a gradual process known as adaptation). If it cannot align with the inoculated fibril, the disease will be blocked.

The following proposals are made about “5 Ts” of the disease phenotype:

1. Tethering: RNA *via* specific tethering, determines protein conformation.
2. Tempo: Palindromic sequences have an advantage, because the primary and complementary chains are identical and can therefore bind to the same protein target. Non-palindromic sequences, in which the primary sequence and complement are different, likely need different protein partners, giving rise to co-pathologies and a slower tempo.
3. Tropism: The ambient pool of small oligonucleotides in particular brain cells offers an advantage to growth and replication of select RNA sequences. Whilst it is envisaged that growth and replication of the agent is able to proceed by recruitment of mononucleotides^9^, particular binucleotides and trinucleotides might also be recruited depending on availability and shape relationships with the protein interface and offer a tempo advantage. This is akin to the cofactor selection hypothesis of neurotropism^34^.
4. Toxicity: Self-annealing repetitive nucleic acids lasso together loops of mitochondrial DNA (mtDNA) into multimeric mtDNA^35,36^, resulting in mitochondrial dysfunction, a known association with neurodegeneration^37^.
5. Transmissibility: Palindromic sequences have a transmissibility advantage (as well as a tempo advantage) because they only need a single protein target.

The proposed arrangement of repetitive RNA braced to protein, suggests repairability, redundancy and protection, potentially confounding claims that RNA is absent from scrapie^7,38^. The arrangement also suggests a structural basis for catalysis, putatively promoting fibril growth and replication^9^.

The name ponc (for **<u>p</u>**rotein **<u>o</u>**rtho-**<u>n</u>**ucleic acid **<u>c</u>**omplex) has been proposed^9^. The ponc hypothesis is testable and has existing experimental support. Poly(A) RNA was added to short basic peptides, including (KL)_3_ and (KL)_5_ *in vitro* and resulted in a complex of RNA and protein in which protein was arranged as amyloid, suggestive of an amyloid nucleic acid (ANA) world, akin to the RNA world^39^. Ponc can be considered a specific structural basis for such an origins of life proposal.

Similarly, RNA and PrP^C^ in the protein misfolding cyclic amplification (PMCA) reaction, form an infectious complex in which RNA is protected^22^. With precautions for biohazard, similar cell-free experiments could be used to test the effectiveness of particular RNAs and proteins in forming novel or known neurodegenerative strains.

Thus, particular peptides and polynucleotides (whether in the primordial soup or today’s brains) may react to form an agent capable of autonomous growth and replication. This shaped-based (analogue) life might have been a step towards sequence-based (digital) life as we know it.

## Methods

### Data sources and selection

Data for this study were sourced from public repositories, the EMDB^6^ and the PDB^11^. Cryo-EM maps and protein models of fibrils from brains of rodents with five scrapie strains were examined: 263K hamster^1^, RML mouse^2^, aRML mouse^3^, a22L mouse^4^ and ME7 mouse^5^. The protein involved in all cases is PrP. In two cases (aRML and a22L) the protein is anchorless. Three strains (263K, aRML and a22L) are from the Rocky Mountain (RM) lab and two (RML and ME7) are from University College London (UCL). This is an important distinction because phosphotungstate (PTA) was used by the UCL lab but not the RM lab in fibril preparation and has a bearing on the interpretation of EDs.

All fibrils had EDs, as determined by inspection and reference to the published papers. Lysine-coordinating EDs and arginine-coordinating EDs were the subject of the present study. EDs from other protein environments (*e.g.* those likely attributable to glycans and GPI anchors) were excluded. EDs were named by the EMDB number and the first coordinating residue. Observations were made on the following cryo-EM maps and atomic models: EMDB 23459 and PDB 7lna in 263K hamster scrapie^1^, EMDB 13989 and PDB 7qig in RML mouse scrapie^2^, EMDB 25824 and PDB 7td6 in aRML mouse scrapie^3^, EMDB 28089 and PDB 8efu in a22L mouse scrapie^4^ and EMDB 15043 and PDB 8a00 in ME7 mouse scrapie^5^. Mouse residue numberings are used in the current paper, unless otherwise specified. Observations were also made on human neurodegenerations for comparison, *viz.* EMDB 26607 and PDB 7umq and EMDB 26613 and PDB 7un5 in PrP fibrils in GSS^27^, EMDB 26663 and 7upe in tau PHF fibrils in AD^33^ and EMDB 10650 and PDB 6xyo in alpha-synuclein fibrils in MSA^17^.

The following were used in the section on candidate constituent molecules: EMDB 25364 and PDB 7sp1 in RNA-induced tau fibrils^15^, PubChem^41^ 90478052 glycogen from oyster, rebuilt as 1,4-linked polyglucose in UCSF ChimeraX^21^, PDB 3iri heparin^42^ and PDB 4l2h poly(ADP-ribose)^43^, extended in UCSF ChimeraX^21^.

### Observations and measurements

Observations and measurements were made in UCSF ChimeraX^21^ unless otherwise stated. Connectivity overlays and some labels were created with ImageJ^44^. A Dell Desktop-GUQ6HDT XPS 15 9500 with Intel® Core™ i7-10750H CPU @2.6 GHz with 16 GB RAM running Windows 11 Home (version 21H2) with broadband internet connection was used to run software and to access data and software online.

Each cryo-EM map was examined at the authors’ recommended contour level with the protein model *in situ*. For clarity, the map was made transparent and clipped to the region of the protein. EDs were identified in their protein environment, by inspection and reference to the published papers.

To determine whether EDs were in perfect register with the protein, markers were placed 30 units apart on EDs and 30 rungs apart on the adjacent protein density and the distances compared. To assess whether EDs were within hydrogen-bonding distance of protein, color zones from the NZ (terminal nitrogen) atoms of lysines were incremented until they reached the surface of the EDs. The angle between 3 markers at 10 unit intervals was measured as an index of straightness.

In some cases, EDs were detached from, or showed minor regions of fusion to, the protein density. In other cases, EDs were extensively fused to the protein density. EDs were isolated from the rest of the density, in order to observe and measure them freely. This was done with the map eraser tool, by erasing the ED and subtracting the remainder from a copy of the original. Where there was extensive fusion between the ED and the protein density, the ED was isolated by using the color zone and split map commands, at a radius of 2 Å from the protein atoms. Further detail was evinced by adjusting the contour level. For some depictions, the hide dust tool was used.

The molecular weight (MW) of the ED was calibrated to the MW of the protein. Protein density was isolated with the color zone and split map commands, with a 2 Å radius around protein atoms. Volumes of isolated EDs and protein densities were measured with the measure blobs tool or measure volume command. The MW of one rung of protein was calculated by opening the protein model in UCSF Chimera^45^, adding hydrogens, selecting one rung and invoking the keyboard shortcut ac mw. The ratio of the volume of ED to the volume of protein density was calculated. The ratio was multiplied by the MW of one rung of protein to give an estimate of the MW of one residue of ED. For comparison, the MW of RNA residues, in Daltons, are as follows: cytosine 304.2, uracil 305.2, adenine 328.2 and guanine 344.2 (average 320.5).

In order to create a molecular model of the protein component of the entire fibril, multiple copies of the PDB protein model were translated and fitted to the corresponding EMDB cryo-EM map using UCSF ChimeraX^21^. Electrostatic potential surfaces were created in UCSF ChimeraX^21^ using the coulombic command.

For comparing the structures of two strains, a script was run in UCSF ChimeraX^21^ in which the root mean square deviation (RMSD) of the alpha-carbon atoms of blocks of 11 residues were calculated sequentially and plotted against the central residue number. For example, the RMSD of residues 95 to 105 was plotted against residue number 100. The chart (created in Microsoft Excel) was annotated with the position of EDs and the variable regions were colour-coordinated with the atomic structures.

ED 28089-147 was split into two near-identical chains after it was observed to appear symmetrical. The ED, viewed at increased level (0.008), was split at a radius of 6 Å (determined empirically) from the terminal nitrogen atoms NH1 and NH2 of R147 in the corresponding protein model PDB 8efu^4^ using the color zone tool in UCSF ChimeraX^21^. The resulting chains were translated vertically in order to view them separately and then one fitted to the other using the fitmap command and the correlation metric noted from the log.

### Molecular docking

A 3mer of oRNA (UUU) was used as ligand. It was chosen with an inter-residue distance as close as possible to the rung distance of the protein. Five rungs of protein (cut down to coordinating and intervening residues) were used as receptor. Ligand and receptor were converted to PDBQT format in AutoDock Tools^46^. Both ligand and receptor were docked as rigid objects. Molecular docking was done with AutoDock Vina^40^. The resulting poses were examined in UCSF ChimeraX^21^. They were assessed for overlap with the ED, symmetrical hydrogen bonds (displayed with relaxed criteria, distance tolerance 0.4 A, angle tolerance 20°) and favourable AutoDock Vina score (affinity, kcal/mol). The protein and selected pose were combined in UCSF Chimera^45^ and submitted to the MolProbity Webserver^47^ to determine the clashscore. The duplex oRNA model was created and examined similarly by designating a 7mer as receptor and 3mer as ligand.

### Ethical statement

The cryo-EM maps and atomic models used in the present study were sourced from the EMDB and PDB public repositories from the publications recorded herein, which for patient data all affirm that ethical review and informed consent were obtained (all patient data for the present study was at one remove and no new patient data was obtained) and for animal data all affirm that appropriate licences were approved and granted (all animal data for the present study was at one remove and no new animal data was obtained).

## Acknowledgements

The present work was done on cryo-EM and atomic models from the EMDB and PDB public repositories. For the human subjects, the patients and relatives were thanked in the original publications and, at one remove, I also thank them. I also thank the scientists, software developers, data managers and funders whose efforts, at one remove, have made my own work possible. Molecular graphics and analyses performed with UCSF Chimera and UCSF ChimeraX were developed by the Resource for Biocomputing, Visualization, and Informatics at the University of California, San Francisco, UCSF Chimera with support from NIH P41-GM103311 and UCSF ChimeraX with support from National Institutes of Health R01-GM129325 and the Office of Cyber Infrastructure and Computational Biology, National Institute of Allergy and Infectious Diseases. I thank St George’s University Hospitals NHS Foundation Trust NHS and South West London Pathology for my employment and St George’s University of London for my honorary academic status.

## Author contribution

LRB performed the whole of this work.

---

**Extended Data Fig. 1.**
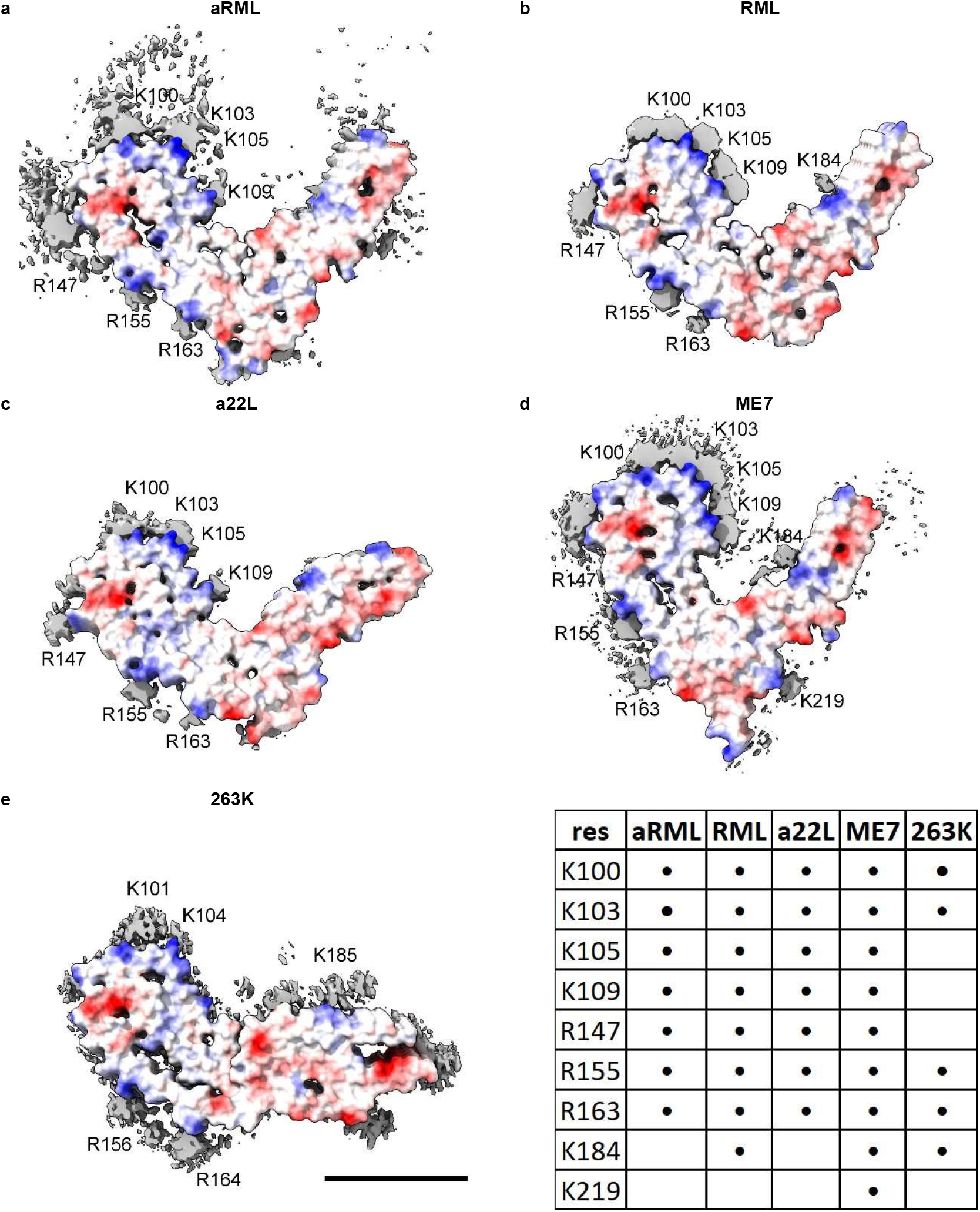
Electrostatic potential surfaces and EDs. Locations of EDs in PrP fibrils from five scrapie strains as summarised in the table. Note lysine patch (K100, K103, K105 and K109) and arginine patch (R147, R155 and R163) on medial and lateral aspects of N-terminal lobe. Electrostatic potential surfaces of proteins are shown, electropositive blue and electronegative red. Images of EMDB 25824 and PDB 7td6 in aRML mouse scrapie^3^ (**a**), 13989 and 7qig in RML mouse scrapie^2^ (**b**), 28089 and 8efu in a22L mouse scrapie^4^ (**c**), 15043 and 8a00 in ME7 mouse scrapie^5^ (**d**) and 23459 and 7lna in 263K hamster scrapie^1^, scale bar 50 Å (**e**), created with UCSF ChimeraX^21^.

**Extended Data Fig. 2.**
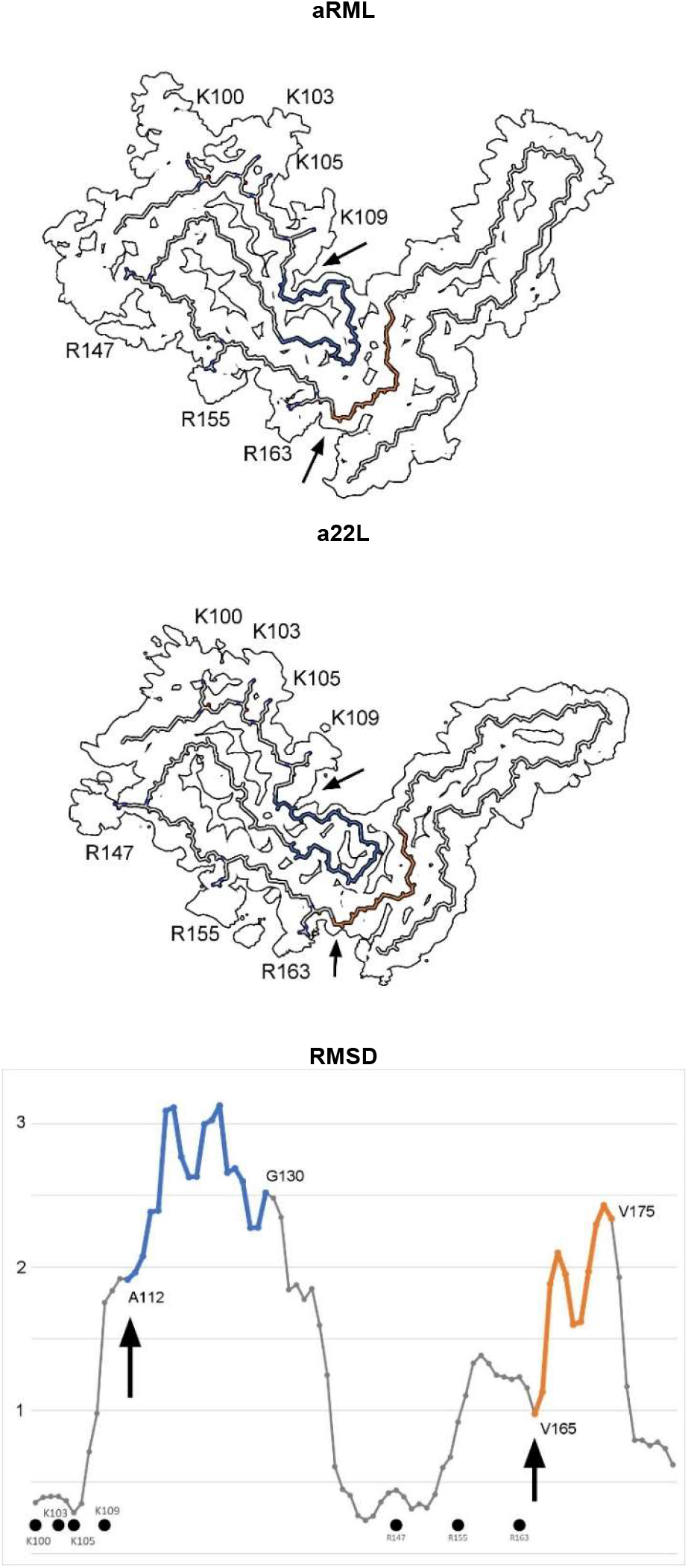
Comparison of scrapie strains. Note difference in angle between N- and C-terminal lobes. The RMSD chart shows peak variability (from arrows) downstream of lysine and arginine patches. The atomic structures indicate articulation at the inter-lobar interface, consistent with an allosteric effect of ligands within EDs at K100, K103, K105, K109, R147, R155 and R163 in syngeneic mice. Images of EMDB 25824 and PDB 7td6 in aRML^3^ (**a**) and 28089 and 8efu in a22L mouse scrapie^4^ (**b**) created with UCSF ChimeraX^21^.

**Extended Data Fig. 3.**
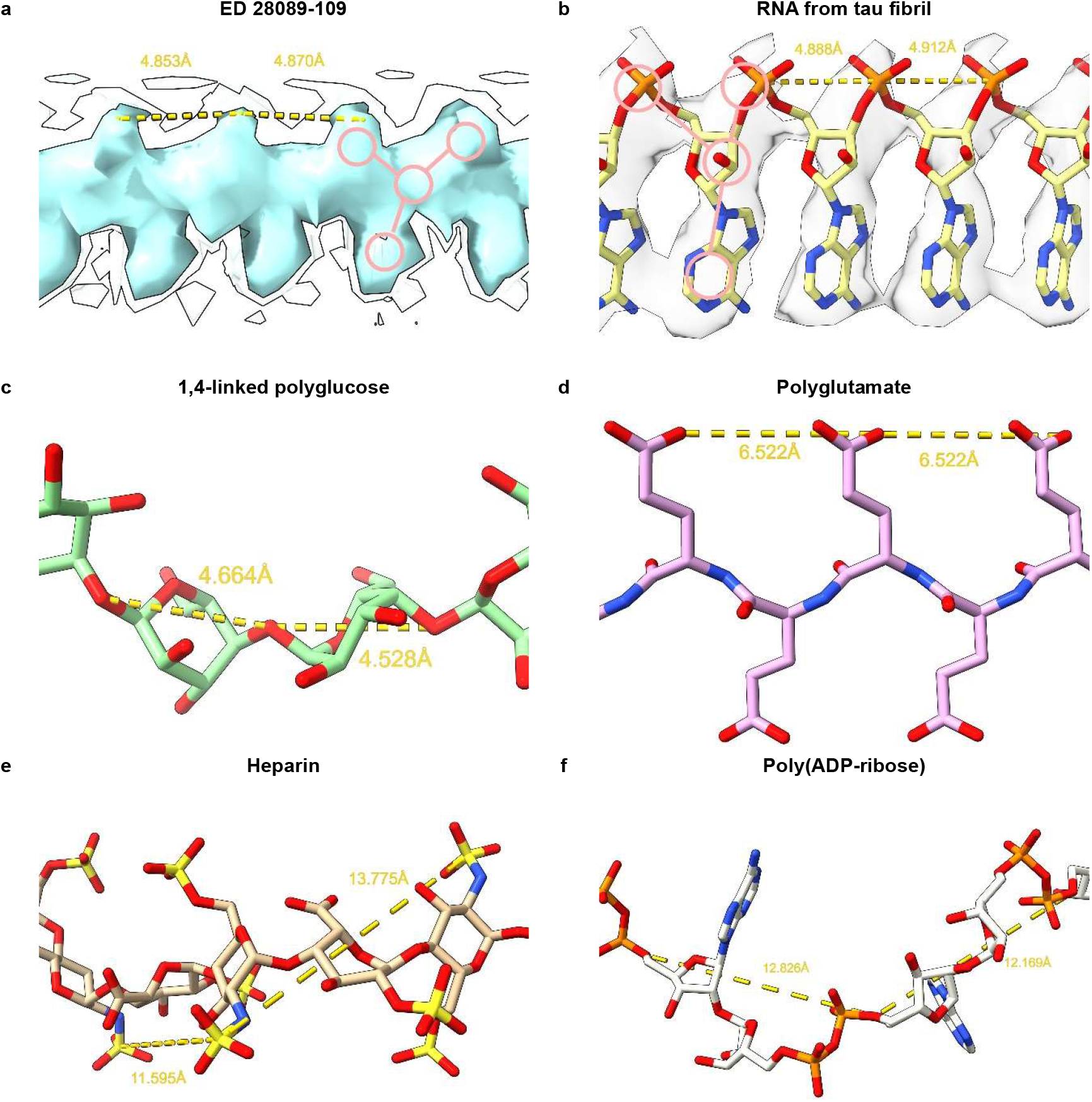
Candidate molecules. (**a**) ED 28089-109 from PrP fibril of a22L mouse scrapie is straight with a 3-blob pattern and Y-shaped connectivity. (**b**) ED from *in vitro* tau fibril, known by experiment to contain a straight form of RNA. Note 3-blob pattern and Y-shaped connectivity. (**c**) 1,4-linked polyglucose is fairly straight with a repeat distance close to that of protein but is not polyanionic and has a single-blob pattern. (**d**) Polyglutamate is a straight polyanion but is multidentate with a repeat distance different to protein. (**e**) Heparin is a coiled, multidentate polyanion with a long repeat distance. (**f**) Poly(ADP-ribose) is a coiled, multidentate polyanion with a long repeat distance. Images of (**a**) EMDB 28089^4^, (**b**) EMDB 25364 and PDB 7sp1^15^, (**c**) Pubchem^41^ 90478052 rebuilt in UCSF ChimeraX^21^, (**d**) polyglutamate built as beta-strand in UCSF ChimeraX^21^, (**e**) PDB 3iri^42^ and (**f**) PDB 4l2h^43^ rebuilt in UCSF ChimeraX^21^, created with UCSF ChimeraX^21^.

**Extended Data Fig. 4.**
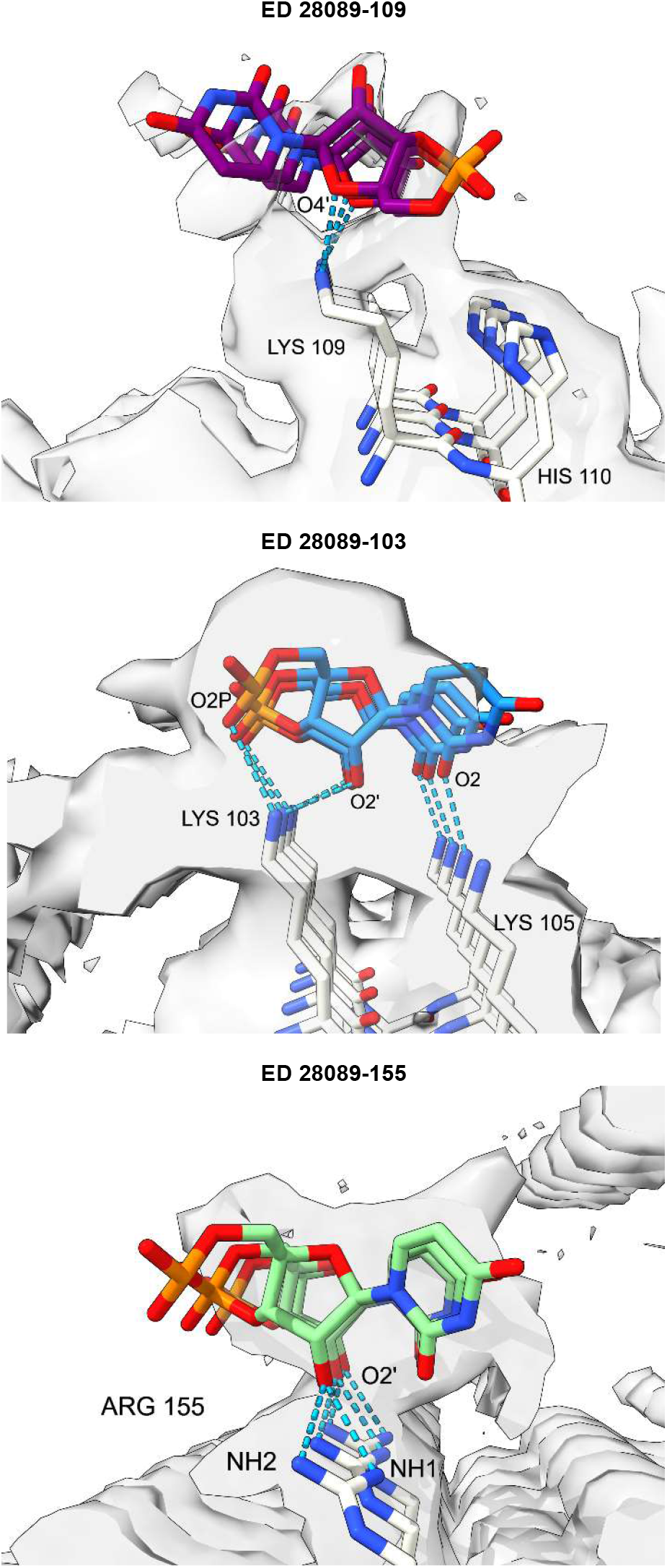
RNA is a feasible ligand for PrP. Ortho-RNA forms a rich symmetrical network of hydrogen bonds with protein, consistent with a role in determining protein conformation. PrP fibril in a22L mouse scrapie. Images of EMDB 28089 and PDB 8efu^4^ created with UCSF Chimerax^21^, oRNA poses generated with AutoDock Vina^40^.

**Extended Data Fig. 5.**
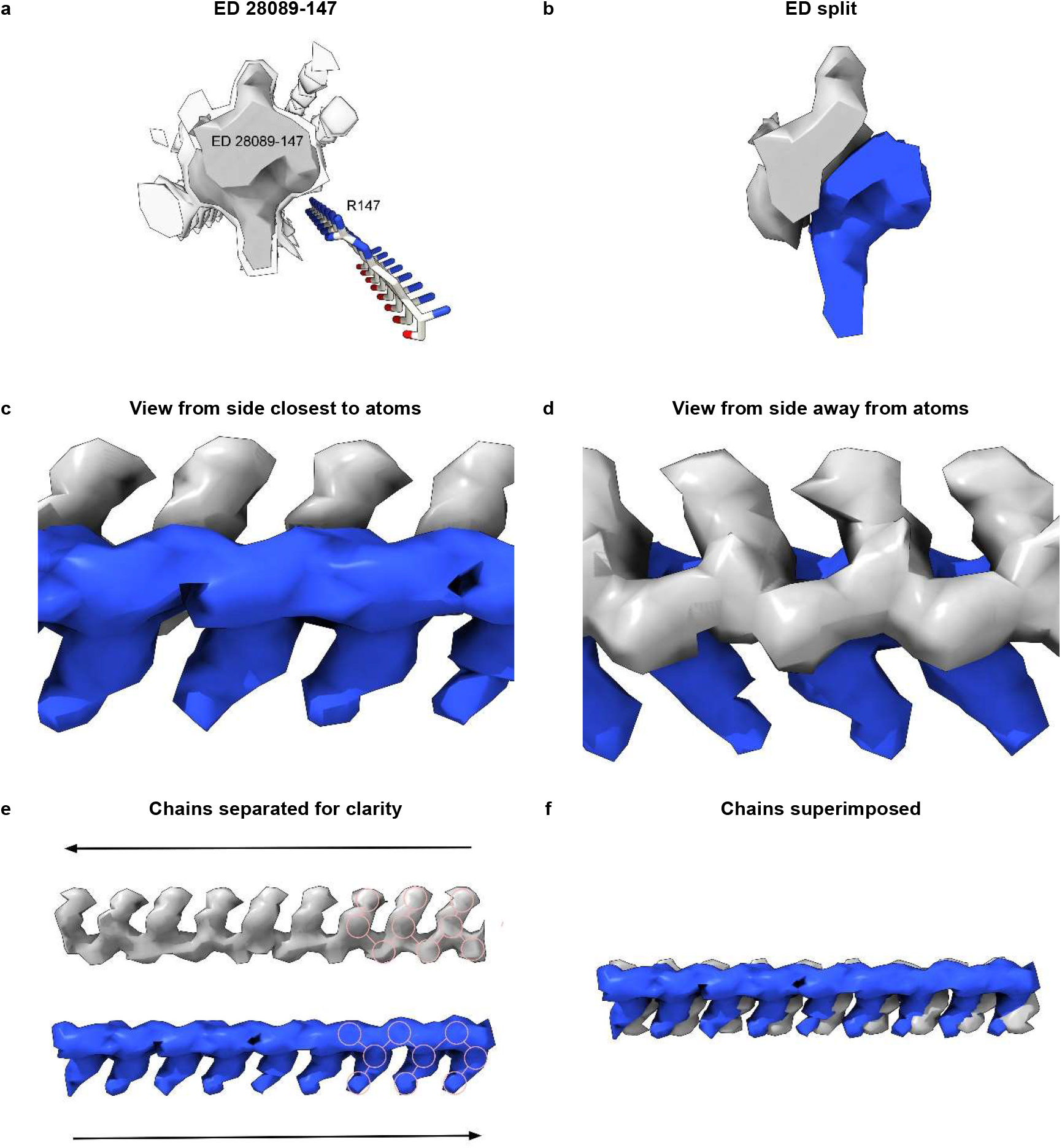
Duplex RNA. (**a**) ED 28089-147 from PrP fibril of a22L mouse scrapie, at authors’ contour level (transparent) and increased level (opaque). (**b**) ED is split by proximity to terminal nitrogens of R147. Note symmetry. (**c**) and (**d**) Side views, indicating two near-identical chains. (**e**) The chains have been separated. Both have a 3-blob pattern and Y-shaped connectivity typical of RNA. Their directions are opposite (anti-parallel). (**f**) The superimposed chains are highly correlated. Images of EMDB 28089 and PDB 8efu^4^ created with UCSF ChimeraX^21^.

**Extended Data Fig. 6.**
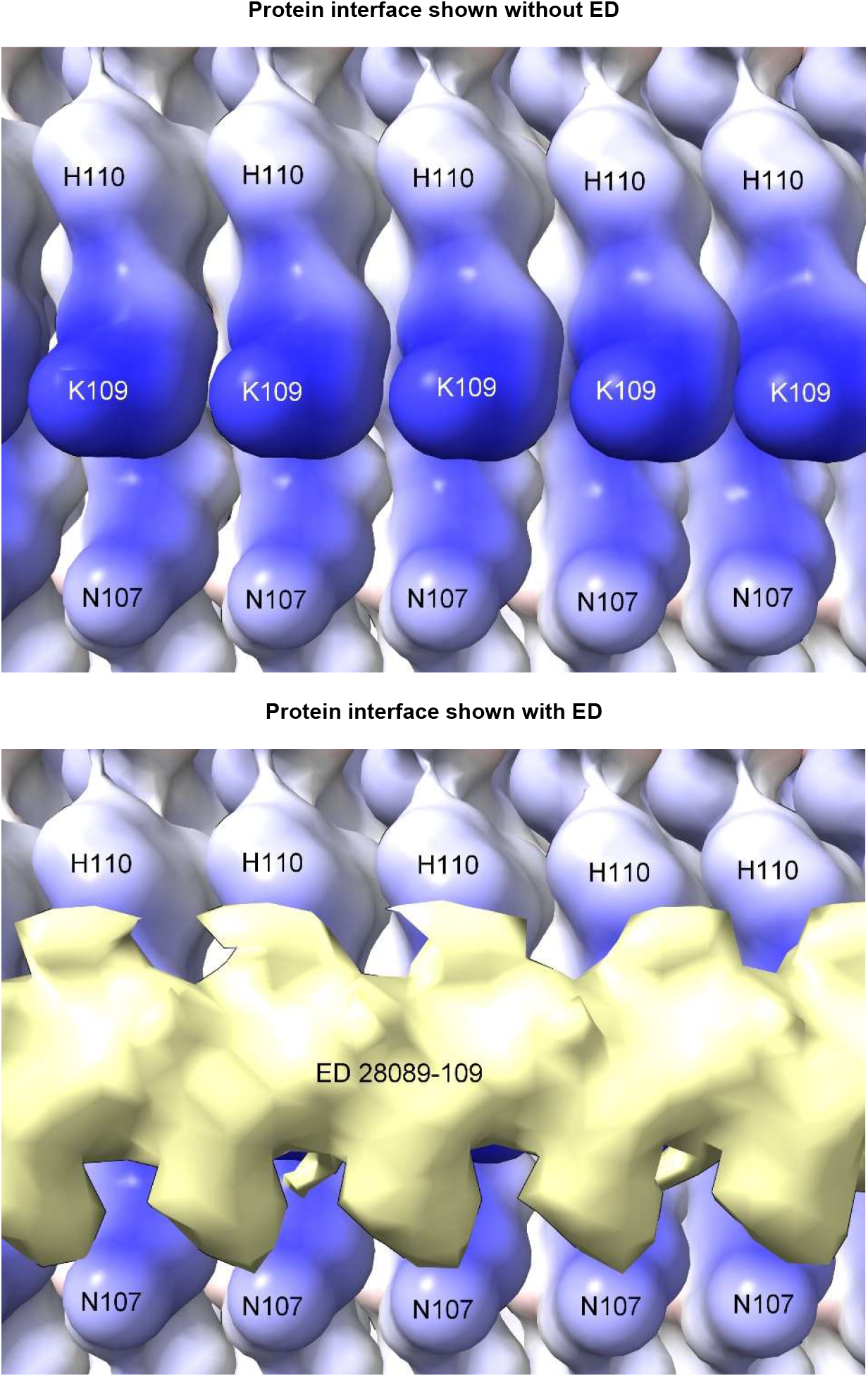
Repetitive interface. PrP fibril from a22L mouse scrapie shown with and without overlying ED. The protein interface is repetitive, suggesting that the constituent molecule of the ED may also be repetitive (*e.g.* a homopolymer or short sequence repeat of RNA). Electrostatic potential surface of protein is shown, electropositive blue and electronegative red. Distance between protein rungs is 4.8 Å. Images of EMDB 28089 and PDB 8efu^4^ created with UCSF ChimeraX^21^.

**Extended Data Fig. 7.**
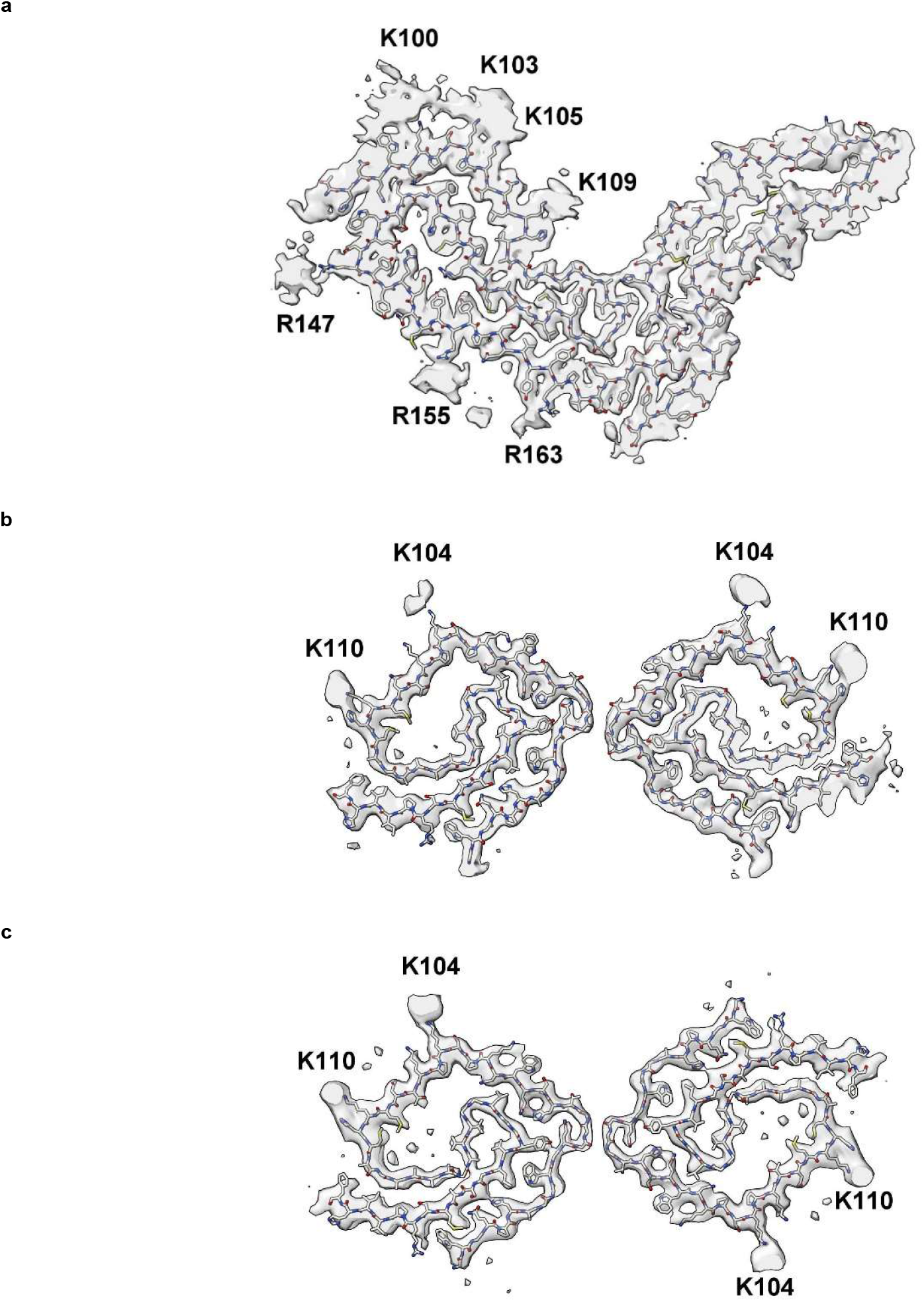
Comparison of scrapie and GSS. PrP fibrils from a22L mouse scrapie (**a**) versus human GSS (**b** and **c**). The scrapie fibril is a single large protofilament whereas the GSS fibrils have two smaller protofilaments. The burden of EDs is greater in scrapie than GSS. Images of EMDB 28089 and PDB 8efu^4^ (**a**), 26613 and 7un5^27^ (**b**) and 26607 and 7umq^27^ (**c**) created with UCSF ChimeraX^21^.

**Extended Data Fig. 8.**
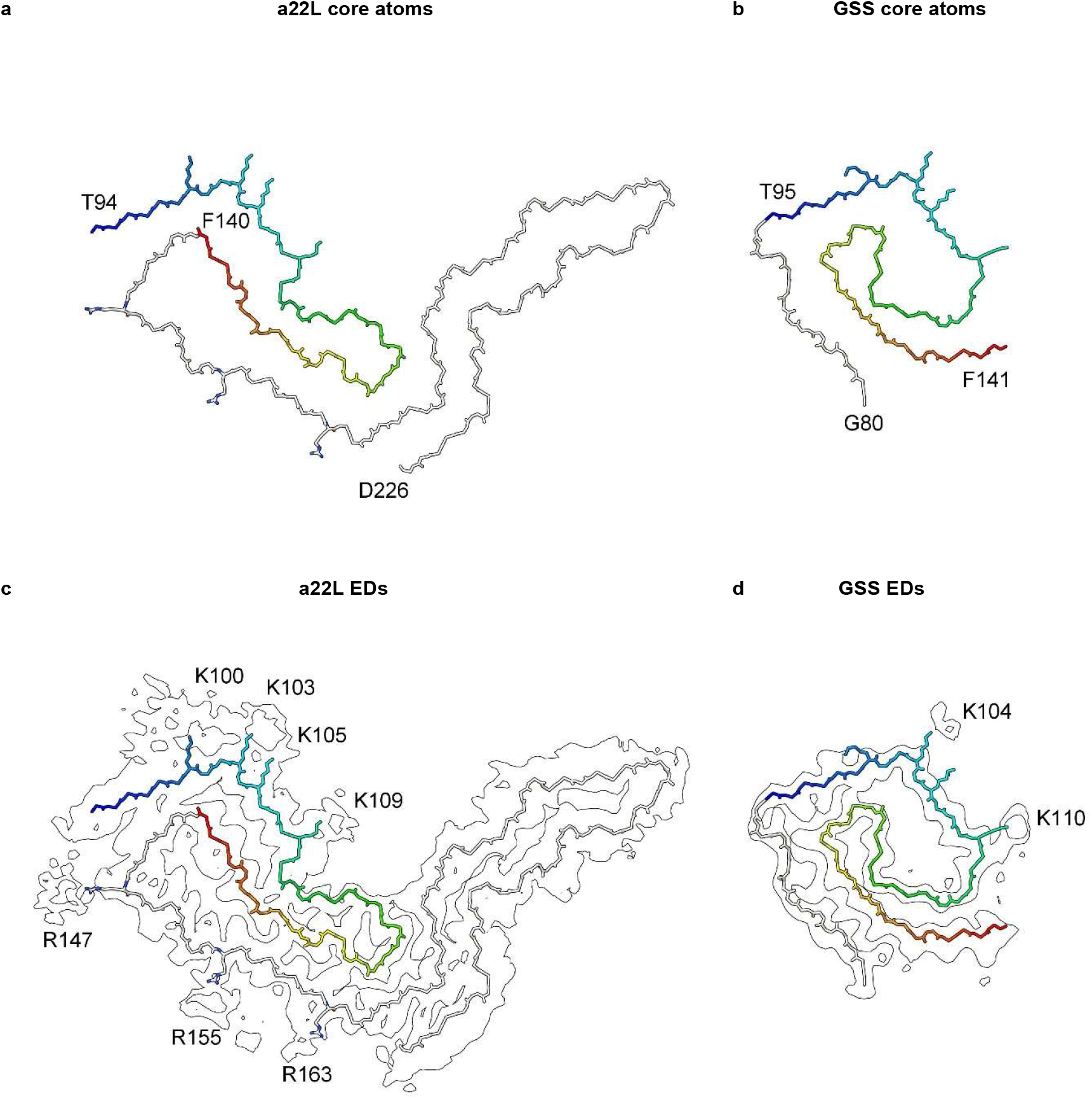
GSS lacks arginine patch. PrP fibrils from a22L mouse scrapie (**a** and **c**) and the human neurodegenerative disease GSS (**b** and **d**). Although the lysine patch is present in both, there are EDs at K100, K103, K105 and K109 in scrapie but only at K104 and K110 in GSS. The arginine patch (with EDs present at R147, R155 and R163 in scrapie) is absent in GSS, due to truncation of the protein core. Images of EMDB 28089 and PDB 8efu^4^ (**a** and **c**) and 26613 and 7un5^27^ (**b** and **d**) created with UCSF ChimeraX^21^.

**Extended Data Fig. 9.**
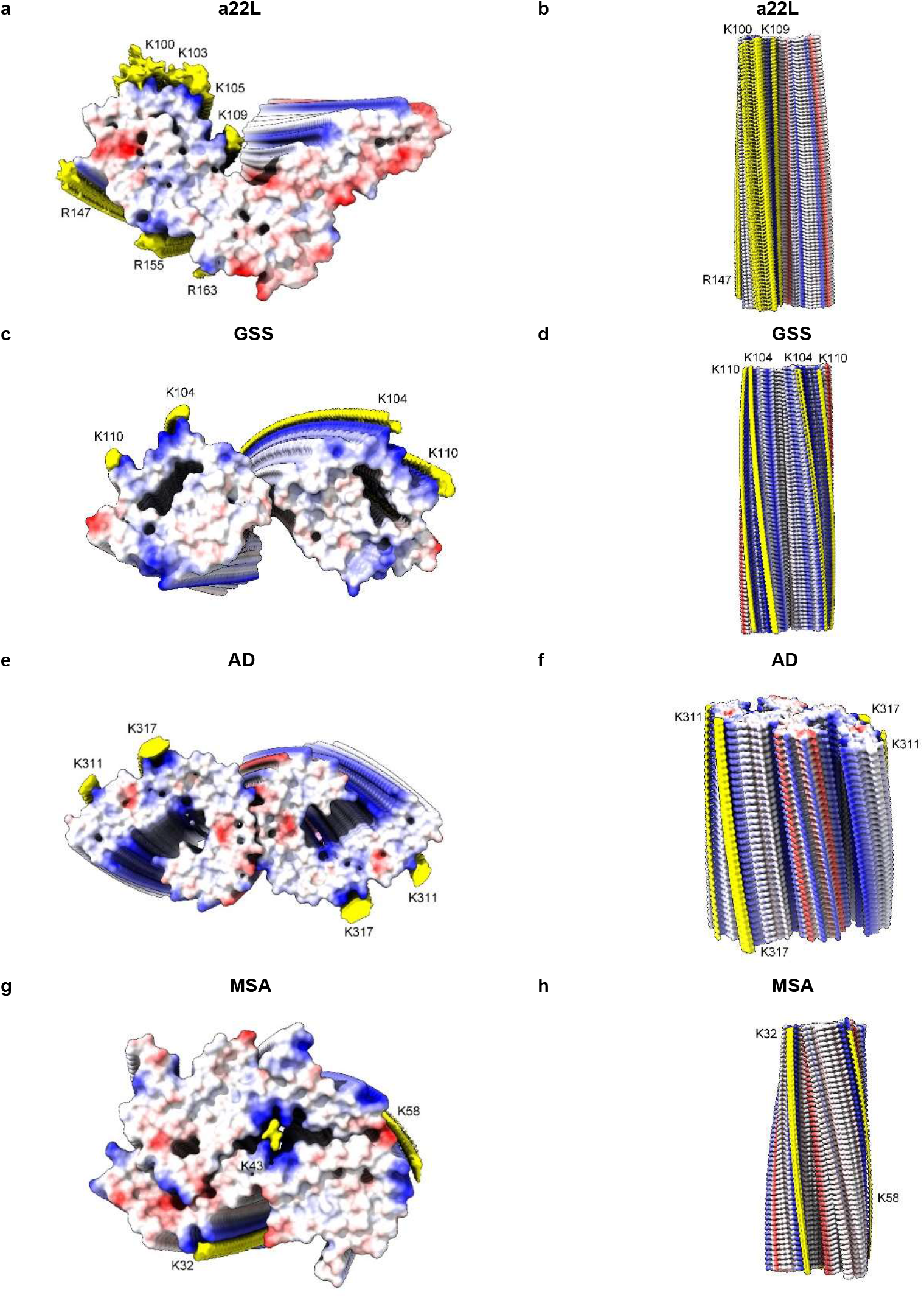
Scrapie and human neurodegenerations. PrP fibrils from a22L mouse scrapie (**a** and **b**) and GSS (**c** and **d**), tau PHF fibril from AD (**e** and **f**) and alpha-synuclein fibril from MSA (**g** and **h**). Electrostatic potential surfaces of proteins are shown, electropositive blue and electronegative red. EDs are shown in yellow. The various fibrils are more similar than different. Distance between protein rungs is about 4.8 Å. Images of EMDB 28089 and PDB 8efu^4^ (**a** and **b**), 26613 and 7un5^27^ (**c** and **d**), 26663 and 7upe^33^ (**e** and **f**) and 10650 and 6xyo^17^ (**g** and **h**) created with UCSF ChimeraX^21^.

